# € Harnessing Protein Language Model for Structure-Based Discovery of Highly Efficient and Robust PET Hydrolases

**DOI:** 10.1101/2024.11.13.623508

**Authors:** Banghao Wu, Bozitao Zhong, Lirong Zheng, Runye Huang, Shifeng Jiang, Mingchen Li, Pan Tan, Liang Hong

## Abstract

Plastic waste, particularly polyethylene terephthalate (PET), presents significant environmental challenges, prompting extensive research into enzymatic biodegradation. Existing PET hydrolases are limited to a narrow sequence space and demonstrate insufficient performance for biodegradation. This study introduces a novel discovery pipeline that combines protein language models (PLMs) with a structural representation tree to identify enzymes based on structural similarity. Using the crystal structure of *Is*PETase as a template, we employed PLMs and a representation tree to efficiently search and cluster target proteins. Screening of candidate proteins was further refined using PLM-based assessments of solubility and thermal stability. Biochemical experiments showed that 14 of 34 candidates exhibited PET degradation activity across a temperature range of 30–60 °C. Notably, we identified a PET hydrolase -*Kb*PETase, which possesses a melting temperature 32 °C higher than that of *Is*PETase and exhibits the highest PET degradation activity within 30–50 °C compared to other wild-type PETases. *Kb*PETase also shows higher catalytic efficiency than FastPETase. X-ray crystallography and molecular dynamics simulations further revealed that *Kb*PETase has a conserved catalytic domain and enhanced intramolecular interactions. This work develops a novel deep learning approach to discover natural PETases with enhanced functions.

## Introduction

Plastic waste poses serious risks to human health and the environment, contributing to pollution, toxic chemical exposure, and ecosystem disruption ^1-3^. Recycling plastic is therefore essential to mitigate these negative impacts, conserve natural resources, and reduce environmental contamination. Polyethylene terephthalate (PET), one of the most widely used plastics, has already been extensively utilized in packaging, textiles, and various consumer products^4^. Due to its durability and extensive use, PET accumulation in landfills and natural ecosystems has become a significant environmental issue ^1,2,5^. Traditional mechanical and chemical recycling methods for PET are limited in both efficiency and environmental sustainability ^6-8^. Consequently, the development of biological solutions for PET degradation, especially using hydrolase enzymes to product high-quality recycled PET (rPET) has obtained significant attentions from both scientific research and industry^9-18^.

Multiple enzymes capable of degrading PET have been biochemically and structurally identified and characterized^19-23^. The most effective PET-degrading enzymes to date are derived from culturable microorganisms isolated from environmental microbial communities, such as the widely used *Ideonella sakaiensis* PET hydrolase (*Is*PETase) ^12^. However, approximately 99% of microbial species in nature are unculturable, making metagenomic data a crucial resource for discovering new enzymes ^24^. Successes in this field include the identification ofleaf and branch compost cutinase (LCC) ^19^, PHL-7 ^22^, Bacterium HR29 (BhrPETase) ^21^ and Thermobifida fusca cutinases (TfCut1 and TfCut2) ^20^ have been reported. Recently, a thermophilic PET-degrading enzyme, MG8, has been discovered from human metagenomic data ^23^. Despite these advancements, the performance of these wild-type (WT) PETases remains insufficient for industrial PET enzymatic hydrolysis. Industrial-scale PET biodegradation requires high-temperature conditions to achieve high yields of rPET ^25,26^, as temperatures approaching PET’s glass transition release molecular energy, facilitating PETase-catalyzed hydrolysis ^18,27-30^. Thus, the major limitations of WT PETases are their low catalytic activities and low thermostabilities ^10,29,31^. Discovering new PET hydrolases with high catalytic activities and thermostabilities are crucial for advancing PETase engineering for the PET biodegradation industry.

Sequence-based screening remains the most widely used approach for enzyme discovery ^32,33^, leading to the identification of enzymes such as LCC and MG8 ^34^. Although sequence-based methods-such as identifying conserved residues, sequence similarities, or hidden Markov models (HMMs)-are effective, there might still remain proteins with similar functional properties that cannot be identifies using these sequence information ^35^. Conversely, because the three-dimensional (3D) structure of a protein largely dictates its function, structure-based discovery becomes a more robust approach and it yields a more diverse repertoire for enzyme mining ^36,37^. While structural data in public databases is still limited ^38,39^, deep learning methods including AlphaFold ^40^ have shown great promise in accurately predicting protein 3D structures, enabling the large-scale exploration and classification of proteins with structure queries.

In this study, we developed a structure-based enzyme discovery pipeline to obtain new PETases with high thermostability and catalytic activity. The pipeline integrates the sequence-based and structure-based retrieval tools, in combinations with protein language model to build a representation tree using sequence embeddings. Based on the predictions, we selected 34 potential thermostable PET hydrolases from two branches of the representation tree for experimental screening. Each candidate protein was expressed and purified, and went through the screening for hydrolytic activity using the substrate analog *p*NPB. Proteins exhibiting degradation activity were further assessed for their ability to degrade amorphous PET film at different temperatures, and their thermostability was characterized by measuring the melting temperature (T_m_). Among the 14 proteins exhibiting PET degradation activity, 8 PETases had a melting temperature (T_m_) more than 10 °C higher than that of *Is*PETase, and 3 proteins demonstrated significantly enhanced degradation activity compared to *Is*PETase.Notably, a PET hydrolase -*Kb*PETaseexhibited a Tm value 32 °C higher than that of *Is*PETase, with superior activity at 30–50 °C compared to other wild-type PETases. Remarkably, its activity at 50 °C is 100 times greater than that of IsPETase and 40% higher than that of LCC at 60 °C.*Kb*PETase also shows enhanced catalytic efficiency (k_cat_/K_M_) compared to FastPETase, one of the most active PETase mutant^41^. The crystal structure of *Kb*PETase revealed a conserved 3D catalytic site, highly similar to the query protein *Is*PETase. Molecular dynamics (MD) simulations revealed that the enhanced intramolecular interactions result in the high thermostability of *Kb*PETase. Overall, this work demonstrates that enzyme mining based on structure and protein language models is a reliable and sensitive approach for identifying and screening candidate PETase, accelerating the discovery of high-performance enzymes.

## Results

### Structure-Base Enzyme Discovery Pipeline

Our structure-based enzyme discovery pipeline for PET hydrolases consists of two primary stages (Fig. 1a). First, we aimed to compile a comprehensive collection of potential PETases sharing structural similarity with known enzymes. Starting with *Is*PETase as the query structure (PDB: 5XFY^42^), we utilized the structure search tool FoldSeek ^43^ to perform structure similarity searches (see details in Methods). Given that structure databases are still limited (e.g., AFDB and ESMAtlas), we employed the identified proteins as queries to conduct an extensive sequence search through the NCBI NR sequence database using MMseqs2 (see details in Methods). This approach results in a vast repertoire of 33,247,501 candidates, which could cover almost all possible PETases that share a similar structure to our query protein, *Is*PETase (Fig. 1b).

**Figure 1.**
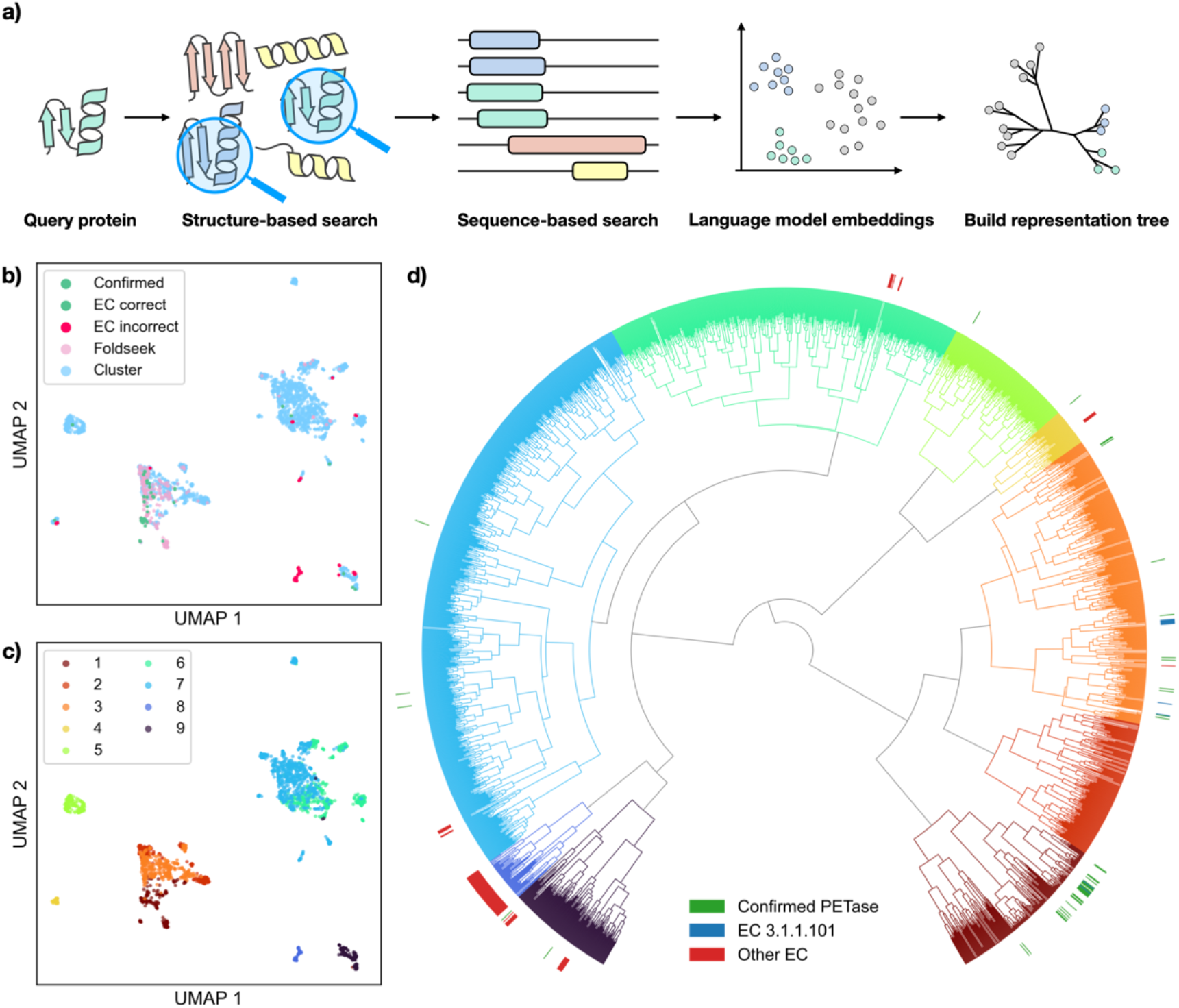
*In silico* structure-based PET hydrolase discovery pipeline and clustering results. (a) The pipeline for PETase discovery combines structure retrieval with FoldSeek, sequence retrieval with MMSeqs2, protein embeddings computation via ProstT5, and representation tree construction to identify functional and robust PET hydrolases. (b) The t-SNE visualization of the PETase candidates identified by ProstT5. Green points indicate proteins with previously validated PET hydrolytic activity, or annotated with corrected EC number 3.1.1.101, while red points represent those not annotated with corresponding EC number. Pink points represent proteins identified using FoldSeek (structure-based search), and orange points correspond to proteins discovered through sequence-based searches. (c) The t-SNE visualization of the PETase candidates colored by representation tree clusters. (d) Representation tree constructed using embeddings from ProstT5. Green bars represent proteins with previously validated PET hydrolytic activity, blue bars are enzymes which with the correct EC number, while the red bars indicate enzymes not annotated with correct EC number. Two clusters are selected for further screening, which are the first cluster (dark red) and the third cluster (orange) reverse clockwise.

The second stage of the pipeline clusters candidate proteins to identify those most likely to exhibit optimal functionality and desired characteristics. Here, we utilized the sequence embeddings from the protein language model ProstT5 ^44^ to represent the proteins. To reduce computational cost, we used representative sequences from 50% identity sequence clusters to calculate embeddings for subsequent analysis. The language model ProstT5^44^ has learned a mapping from protein sequences to structures, which effectively represent the proteins in the structure space. This means that proteins with closer embeddings in Euclidean space and are more likely to share similar structures. Subsequently, agglomerative clustering was applied to the protein embeddings to group the candidate enzymes and build representation tree (Fig 1c and d). To elucidate the functions and characteristics of each cluster, a list of well-studied enzymes with annotations are clustered together for comparison. The agglomerative clusters enabled us to pinpoint the most promising clade for PET degradation activity which has more confirmed PET hydrolases, while others with enzymes from other EC numbers (Fig. 1d). For further screening, we selected two clusters and predicted the melting temperature (T_m_) and solubility for the candidates within these clusters using a fine-tuned ESM2 ^45,46^ model and ProtSolM ^47^, respectively. Finally, 34 candidates were selected for wet-lab experiments after AlphaFold2^48^ structure prediction and filtration.

### Characterizations of Putative PET Hydrolases

Figure 2a illustrates our biochemical characterization workflow. We first analyzed the sequences of 34 candidates to identify signal peptides (Supplementary Fig. 2). SignalP was used to detect these peptides, and truncation was applied to improve expression and purification efficiency ^49,50^. Following codon optimization for enhanced expression (Supplementary Table 1), the modified sequences were cloned into the pET28a (+) vector. The constructs were then expressed in *E. coli* BL21 (DE3), employing an N-terminal hexa-histidine tag to facilitate purification. Using this biochemical method, 26 of the 34 PETase candidates were successfully expressed and purified.The functions of the expressed PETase candidates were subsequently tested. T For rapid evaluation of PET degradation, *p*-nitrophenyl butyrate (*p*NPB) was used as a substrate. PETase catalyzes the hydrolysis of ester bonds in *p*NPB, yielding *p*-nitrophenol and butyric acid. The p-nitrophenol exhibits a strong absorbance peak at 410 nm, which reflects the activity of proteins by calculating the peak area ^51^ (Supplementary Fig. 3). After incubating proteins with substrates at 37 °C, 14 of the 26 expressible proteins exhibited ester bond-cleaving activity. (Fig. 2b, Supplementary Fig. 4). This result suggests that our pipeline has high success rates in finding PETase with ester bond cleavage functions.

**Figure 2.**
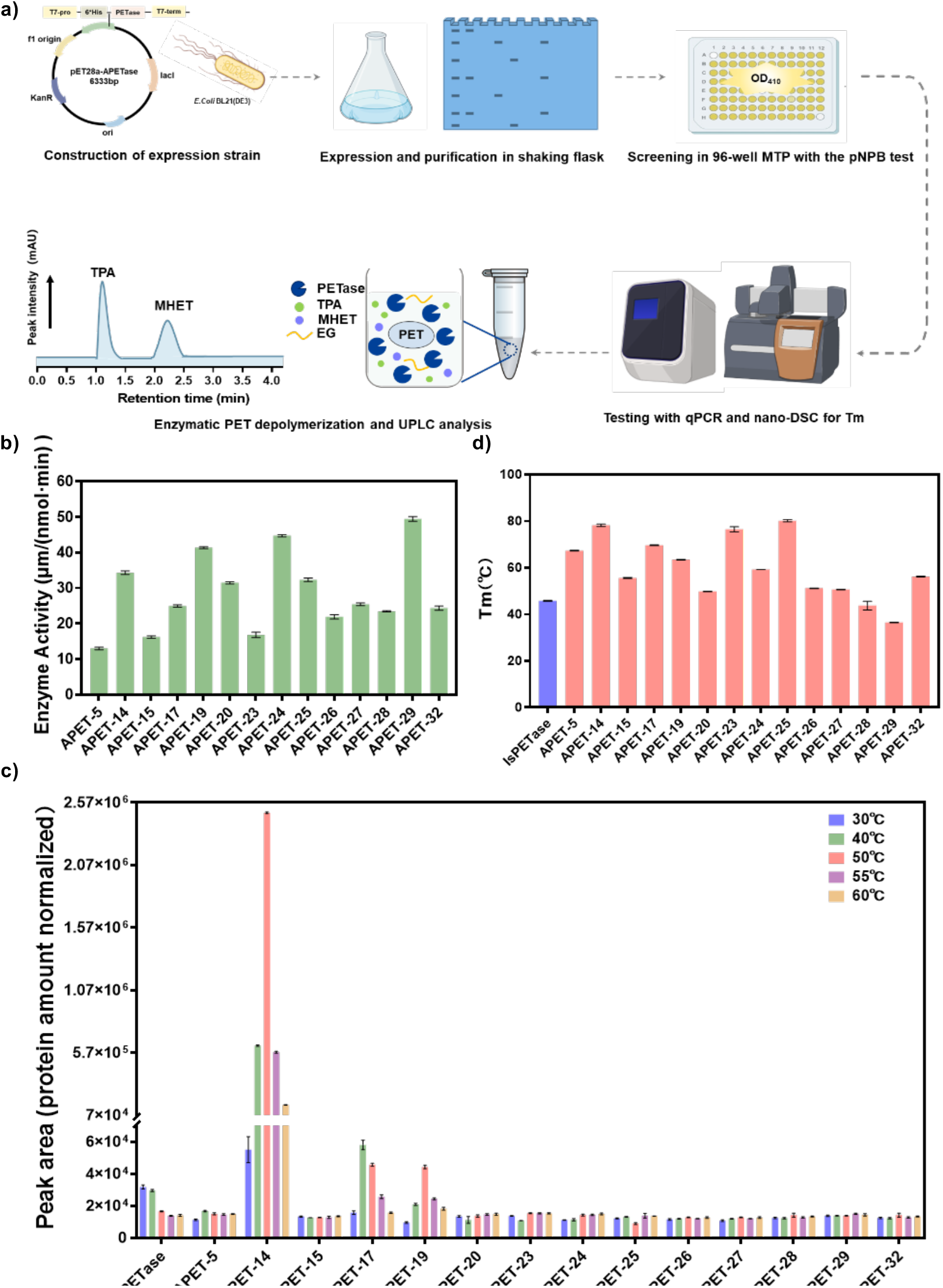
Experimental validation and biochemical characterization of candidates. (a) Experimental validation workflow for the discovered PETases. (b) Relative enzyme activity of 14 proteins showing effective ester bond hydrolysis, as measured using *p*NPB as a substrate at 37 °C. Reactions were performed in triplicate; Data are presented as mean values ± SD. (c) Compare the degradation activity of candidate proteins on amorphous PET films. The reaction was conducted in 50 mM Glycine-NaOH (pH 9.0) at various temperatures (30 °C, 40 °C, 50 °C, 55 °C and 60 °C) for 72 h. Reactions were performed in triplicate; Data are presented as mean values ± SD. (d) The Tm values of the 14 proteins exhibiting ester bond cleavage activity. Reactions were performed in triplicate; Data are presented as mean values ± SD.

To evaluate the PET degradation performance of proteins with ester bond cleavage activity, we utilized a semi-quantitative screening approach^49^. We utilized commercially available amorphous PET film (AF-PET, Goodfellow Cambridge Ltd, Cat. No. ES301445) as the substrate, widely used for assessing PET degradation activity ^13,41^. The AF-PET was uniformly cut into circular discs with a diameter of 6 mm for each reaction. The aromatic reaction products mono(2-hydroxyethyl) terephthalate (MHET) and terephthalic acid (TPA) were quantified using ultra-high-performance liquid chromatography (UPLC)^13^. To enable comparisons with the previously reported PET hydrolases, we used *Is*PETase, the reference protein during structural searches, as a control baseline. Since the optimal catalytic temperatures of the enzymes were initially unknown, we tested the PET degradation activity of the 14 candidates across a range of 30-60 °C. All proteins exhibited activity within this range, with 11 of 14 displaying degradation activity comparable to *Is*PETase. Among these, APET-14 showed the highest catalytic activity at 50 °C, with a 100-fold increase compared to *Is*PETase (Fig. 2c). Given that melting temperature (Tm) is an indicator of PETase thermostability and performance ^11^,we used differential scanning fluorimetry (DSF) to determine the T_m._ of the proteins. The results revealed that the T_m_ values of the 14 proteins ranged from 36.4 °C to 80.1 °C (Fig. 2d), indicating the pipeline’s effectiveness in identifying thermostable, functional candidates.

Notably, APET-14 exhibited excellent thermostability, exhibiting a Tm of 78.2 °C, which represents a 32.4 °C increase compared to *Is*PETase. This result was confirmed by nanoDSC analysis, consistent with DSF data (Supplementary Fig. 5). These findings suggested that APET-14 possesses the traits of an efficient PET-degrading enzyme, exhibiting both high enzymatic activity and thermostability. Therefore, we selected APET-14 - *Kb*PETase for further evaluation. Furthermore, the sequence similarity between *Kb*PETase and known PETases currently ranges from 30% to 50%, highlighting the capability of our pipeline to identify functional hydrolytic proteins in regions of low sequence similarity. (Supplementary Fig. 1).

### Identification of *Kb*PETase exhibiting high PET hydrolytic activity and thermostability

We compared the activity of *Kb*PETase to that of other characterized PET degrading enzymes, such as *Is*PETase, PE-H, BAT, Cut_190, Thc_Cut1, TFH, and LCC, using AF-PET as the substrate across a temperature range of 30-60 °C (Fig. 3a) Mesophilic proteins such as *Is*PETase, PE-H and BTA-2 were included for comparative analysis ^12,52,53^, alongside thermophilic proteins LCC and TFH, to ensure a broad temperature range of activity. ^19,54^. Furthermore, Cut190, *Is*PETase, LCC, and PE-H are recognized as superior starting templates for contemporary directed evolution initiatives ^18^.

**Figure 3.**
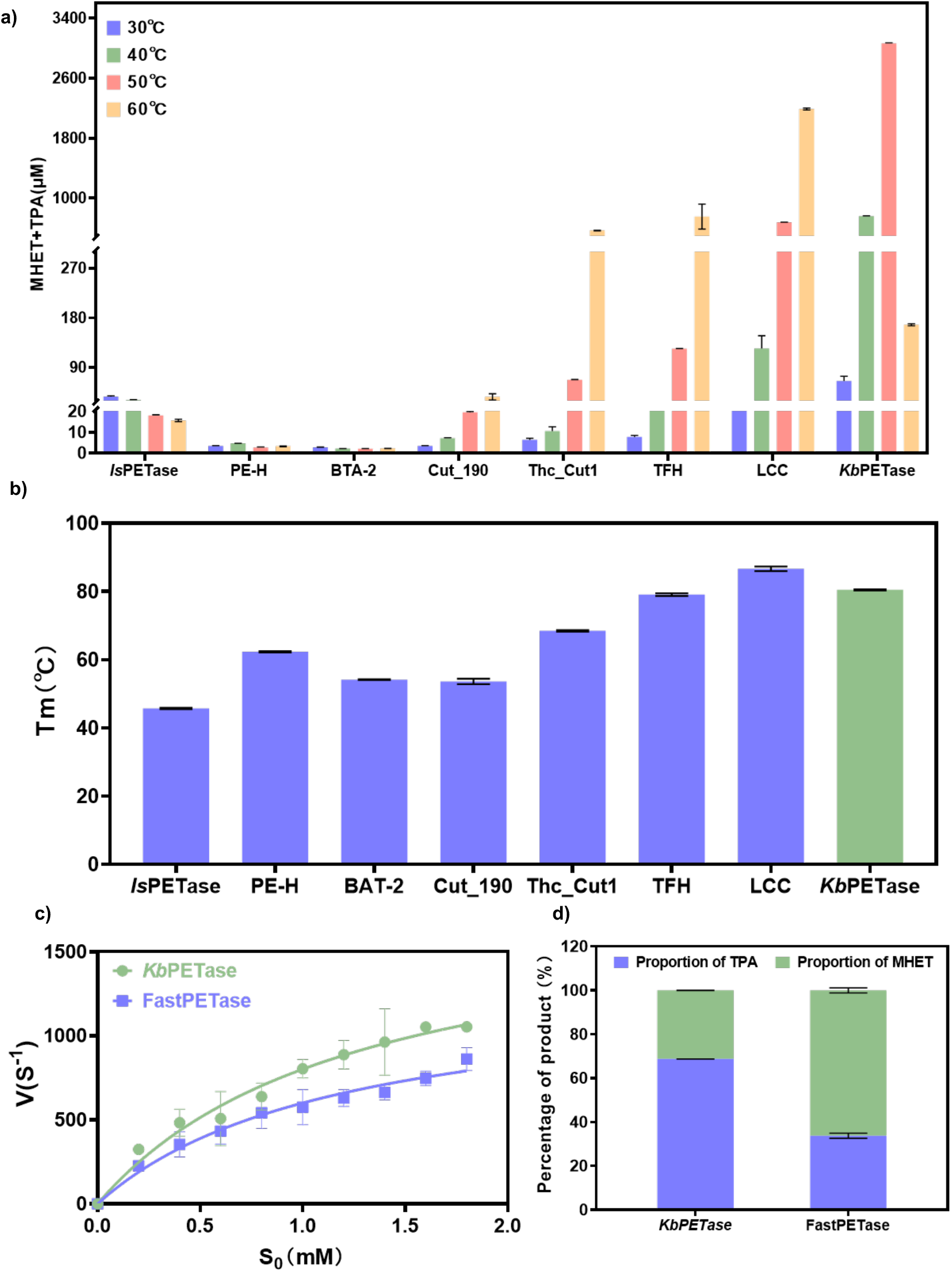
Activity and thermostability measurements of discovered PETases (a) PET film degradation activity of the previously characterized PETases compared to *Kb*PETase. The reaction was conducted in 50 mM Glycine-NaOH (pH 9.0) at various temperatures (30 °C, 40 °C, 50 °C, 55 °C and 60 °C) for 72 h. Reactions were performed in triplicate; Data are presented as mean values ± SD. (b) Comparison of the Melting temperatures of *Kb*PETase with other reported wild-type PETases using DSF. Reactions were performed in triplicate; Data are presented as mean values ± SD. (c) Comparison of the enzymatic kinetics curves of *Kb*PETase and FastPETase at 50°C using p-nitrophenyl butyrate (*p*NPB) as the substrate. Reactions were performed in triplicate; Data are presented as mean values ± SD. (d) The percentages of PET depolymerization were determined by analyzing the products released during the process at 50C°, as measured through UPLC. Reactions were performed in triplicate; Data are presented as mean values ± SD.

*Kb*PETase exhibited high PET degradation activity across temperatures of 30, 40, and 50 °C. In the reaction at 30 °C, *Kb*PETase outperformed both LCC and TFH, displaying double the activity of *Is*PETase, which is the most effective wild-type PETase at moderate temperatures^12^ (Fig. 3a). At 40 °C, the degradation activity of *Kb*PETase was 20 times greater than that of *Is*PETase, and at 50 °C, it was 100 times higher (Fig. 3a). To assess thermostability, we analyzed the melting temperatures of *Kb*PETase and other enzymes, finding that KbPETase’s enhanced activity likely correlates with its superior thermostability (Fig. 3b). When assessing the degradation performance at 50°C, *Kb*PETase exhibited a significantly higher activity, showing five times the enzymatic activity of LCC under the same temperature conditions (Fig. 3a). Furthermore, *<u>Kb</u>*PETase demonstrated enhanced activity relative to other thermophilic enzymes, with its degradation efficiency at 50°C exceeding LCC at 60°C by 40%, despite the trend for higher temperatures to favor PET degradation^7,17,18^ (Fig. 3a). Despite the melting temperature is the second highest among the tested enzymes, the activity of *Kb*PETase at this elevated temperature was still inferior to that of TFH and LCC, both of which possess similar T_m_ values (Fig. 3b). These findings indicate that *Kb*PETase combines the moderate temperature catalytic range typical of mesophilic enzymes with the elevated activity seen in thermophilic counterparts.

We also compared *Kb*PETase with FastPETase, the most active mutant derived from *Is*PETase known for its optimal catalytic performance at 50 °C ^41^. Using *p*NPB as substrate, we evaluated the Michaelis-Menten kinetic parameters (Fig. 3c). As shown in Table 1, *Kb*PETase exhibited a catalytic constant (K_cat_) 1.4 times higher than that of FastPETase, with no significant changes in K_m_. Furthermore, upon assessing K_cat_/K_m_, *Kb*PETase demonstrated a catalytic efficiency that was 1.3 times greater than that of FastPETase, indicating that its elevated catalytic activity may indeed stem from its enhanced catalytic efficiency. The proportion of terephthalic acid (TPA) among the degradation products of PET is crucial for PET recycling ^25,55^. Notably, *Kb*PETase produces TPA at a concentration that is three times higher than that of FastPETase (Fig. 3d). These results suggestthat *Kb*PETase could serve as a more promising template for enzyme engineering compared to other WT enzymes exhibiting extreme mesophilic or thermophilic traits, such as *Is*PETase and LCC.

**Table 1.** Kinetic parameters of *Kb*PETas and FastPETase derived from the Michaelis–Menten experiments.

| | $K_m$ (mM) | $K_{cat}$ (S<sup>-1</sup>) | $K_{cat}/K_m$ (mM<sup>-1</sup>·S<sup>-1</sup>) |
| --- | --- | --- | --- |
| ***KbPETase*** | 1.275 | 267.373 | 209.70 |
| ***FastPETase*** | 1.187 | 192.427 | 162.11 |

### Sequence and structure analysis of *Kb*PETase

To provide structural insights into the enhanced PET-degrading activity of *Kb*PETase, we determined its crystal structure using X-ray crystallography at a resolution of 1.75 Å (PDB: 9IW9). Overall, the structure of *Kb*PETase exhibits a high degree of conservation when compared to *Is*PETase (PDB: 5XH3) (Fig. 4a). The catalytic triad (S128-H206-D176) and substrate pocket of *Kb*PETase are also largely conserved compared to those of *Is*PETase (S131-H208-D177) (Fig. 4b and c). This conservation indicates that our protein mining strategy not only retains the overall structure scaffold but also preserves the active site, ensuring the enzymatic activity necessary for effective PET degradation.

**Figure 4.**
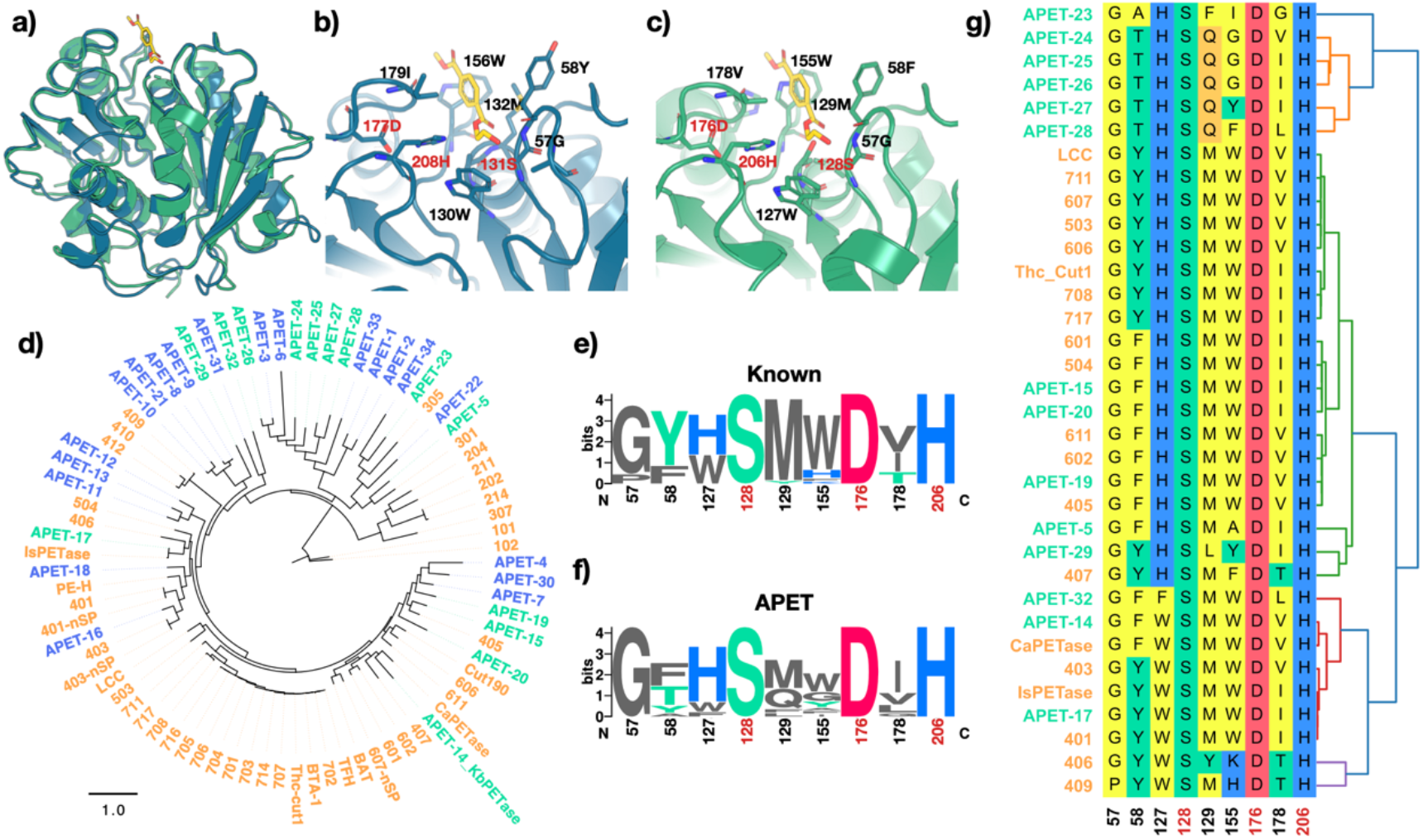
Structure and sequence characteristics of APET enzymes and known PETase. (a) X-ray structure of *Kb*PETase (green, PDB: 9IW9) and *Is*PETase (blue, PDB: 5XH3). The substrate structure (yellow) is from *Is*PETase structure and structural aligned to *Kb*PETase pocket. (b,c) Pocket structure of *Is*PETase (b) and *Kb*PETase (c). (d) Phylogenetic tree of the 34 candidate proteins selected for experimental validation (blue and green), alongside previously reported PET-degrading enzymes (orange). The APET candidates with confirmed PET hydrolase activity are marked with green, otherwise in blue. (e, f) The sequence logo plot for the pocket residues. Height indicates the conservation score for each site, indexed by the location in *Kb*PETase. (g) Structure-based alignment result for pocket residues from APET (green text) and known PETases (orange text). Residues are marked by their biochemical properties, and hierarchical clustering presented on the right. Residue indices are from *Kb*PETase.

A phylogenetic tree was constructed to include APET candidates alongside known PET hydrolases (Fig. 4d). The discovered enzymes are broadly distributed throughout the tree. One major clade contains most known PETases, including *Is*PETase, LCC, Cut190, and Thc_cut1, from which *Kb*PETase also originates. Furthermore, several new clades emerged, populated by APET candidates positioned closer to the root of the phylogenetic tree. This arrangement suggests that the structure-based discovery pipeline may effectively identify ancestral enzymes based on conserved structural features.

To further investigate the substrate pocket, structure alignment was performed using TM-align. Compared with reported PETases, the newly discovered APETs exhibit greater diversity in pocket residues while retaining a conserved catalytic triad (Fig. 4e– g). Notably, residues at positions 58, 129, and 155 show higher variability compared to known PET hydrolases (Fig. 4e and f). Hierarchical clustering based on pocket residues also revealed that APETs are distributed across all major types of known PET hydrolases.

### Molecular dynamics analysis of *Kb*PETase

To further understand the underlying factors for enhanced thermostability of *Kb*PETase, we performed all-atom MD simulations comparing *Kb*PETase with *Is*PETase. We used *Is*PETase because it serves as the initial template enzyme during the search process and acts as a representative of mesophilic PETase. As shown in Figure 5a, the root mean square fluctuation (RMSF) data for the proteins revealed that *Kb*PETase exhibits lower conformational dynamics and reduced flexibility compared to *Is*PETase. The flexibility of proteins is greatly decreased when stronger intramolecular interactions is present. The increased rigidity of *Kb*PETase is further verified in Figure 5d and e, wherein a comparative analysis of the conformational phase space sampled during MD simulations for both *Kb*PETase and *Is*PETase was undertaken. Notably, the majority of *Kb*PETase conformations were identified at a reduced radius of gyration and diminished RMSD in comparison to *Is*PETase. This observation underscores the presence of stronger intramolecular interactions within *Kb*PETase throughout the MD process, conferring a more compact overall structural organization. Remarkably, *Kb*PETase encompasses a greater number of hydrogen bonds than *Is*PETase across its entire molecular structure, despite both enzymes sharing an equivalent count of salt bridges (Figure 5b). Furthermore, an increased presence of hydrogen bonds within the catalytic center of *Kb*PETase was recorded relative to *Is*PETase (Figure 5c). These enhanced intramolecular interactions significantly increase the rigidity of the protein structure, thereby enhancing its overall thermostability. This result is in agreement with the observation that T_m_ of *Kb*PETase is higher than *Is*PETase, which indicates that the high activity of *Kb*PETase at ambient temperature is influenced not only by its high catalytic efficiency (Table 1) but also by its enhanced stability. These findings suggest that limited conformational mobility may contribute to its enhanced thermostability, making it a promising candidate for PET biodegradation across a range of temperatures.

**Figure 5.**
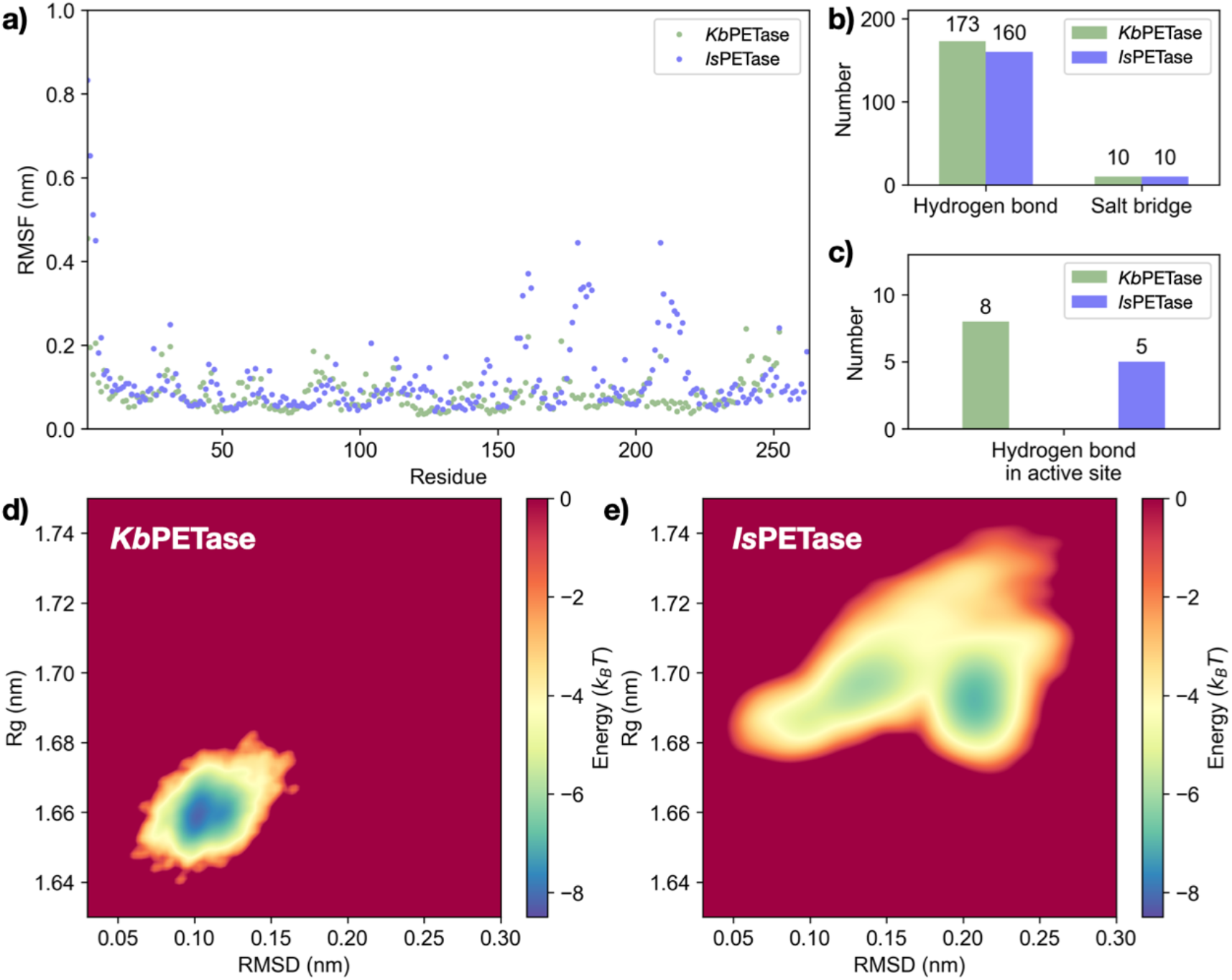
Dynamics analysis of *Kb*PETase and *Is*PETase. (a) Comparison of root mean squared fluctuations (RMSF) of *Kb*PETase and *Is*PETase, depicted over the course of 100 ns production simulations. (b) The global number of hydrogen bonds and salt bridges in *Kb*PETase and *Is*PETase.(c) The localized number of hydrogen bonds and salt bridges around the catalytic sites in both enzymes. (d) and (e) Free energy landscape of *Kb*PETase and *Is*PETase, where RMSD (root mean squared deviation) reflects the structural deviations during production simulations relative to the initial MD snapshot post-equilibration. The free energy indicates the relative frequency of specific conformations with given RMSD and radius of gyration (*R*g) throughout the MD process.

## Discussion and Conclusion

The increasing environmental impact of plastic waste, particularly PET, underscores the urgent need for innovative and effective biodegradation strategies ^5,8,11,26^. Consequently, the search for promising PET-degrading enzymes has been widely pursued, leading to the successful characterization of several proteins to date ^12,18-21,23^. However, current enzyme discovery methods heavily rely on sequence-based analysis, lacking effective structural screening tools to broaden the exploration space. This reliance limits PETase discovery to a narrow sequence landscape ^49^. In this study, we developed a structure-based enzyme discovery pipeline to obtain novel PETases with high thermostability and catalytic activity, by using FoldSeek for structural similarities with *Is*PETase. Through sequence alignment with the NCBI database via MMseqs, we generated a diverse pool of candidate enzymes, further refined by hierarchical clustering using the protein language model, ProtT5, to select the most promising candidates for PET degradation. We utilized the protein language model ProstT5 to perform hierarchical clustering, resulting in a refined list of candidates with potential PET degradation capabilities. The t-SNE visualization demonstrated an expanded search space achieved through this multi-step approach, underscoring the diversity of potential candidates.The application of ESM-2 for T_m_ predictions and ProtSolM for solubility assessments ensured that only candidates with superior properties relative to *Is*PETase were advanced. The sequence identity between our 34 candidate enzymes and known wild-type PETases consistently remains below 40%, which enables the discovery of a substantial number of proteins with low sequence similarity, broadening the potential for enzyme engineering.

Subsequent activity and thermostability screenings identified 14 candidates with PET-degrading capability, among which 10 exhibited T_m_ values higher than *Is*PETase, and 11 demonstrated comparable degradation activity. These results indicate that structural screening effectively enhances the identification of functionally relevant proteins, and that PLMs facilitate a more efficient selection process. Among the candidates, we identified *Kb*PETase, a PET-degrading enzyme that combines characteristics of both mesophilic and thermophilic enzymes, enabling it to catalyze efficiently at moderate temperatures while exhibiting high catalytic activity and a T_m_ typical of thermophiles. *Kb*PETase demonstrated a T_m_ increase of 32.4 °C relative to *Is*PETase, though lower than LCC. Additionally, *Kb*PETase exhibited remarkable PET degradation at ambient temperature, with hydrolytic activity at 50 °C reaching 100-fold that of *Is*PETase, surpassing the degradation performance of LCC at 60 °C by 40%. When compared to FastPETase, the most active variant of *Is*PETase, *Kb*PETase exhibits superior catalytic efficiency and lower TPA production, which is crucial for efficient PET recycling^25,55^. Due to its high catalytic efficiency, superior PET degradation activity, and excellent thermal stability, along with its ability to catalyze the production of a high proportion of TPA, *Kb*PETase shows great promise as an ideal candidate for subsequent directed evolution. This makes it a valuable starting protein for advancing bioremediation strategies, offering a new choice for PET biodegradation.

The crystal structure of *Kb*PETase revealed a highly conserved catalytic domain and pocket residues when compared to *Is*PETase and other known PETases, underpinning the enzyme’s PET degradation efficacy. MD simulations further confirmed that *Kb*PETase possesses more intramolecular interactions, a more stable conformation, and a lower energy state, all contributing to its enhanced thermostability. The structure-function relationship elucidated in this study highlights the crucial role of specific residues in substrate binding and catalysis, confirming the success of our mining strategy in not only identifying novel enzymes but also in preserving the functional integrity of their active sites.

Our pipeline not only expands the repertoire of known PET hydrolases but also establishes a robust framework for future enzyme discovery. With the advancements in deep learning and protein language models, it may soon be possible to predict protein structures, and even complexes, more accurately, enabling the preliminary selection of candidate proteins through modeling. This approach not only aids in identifying exceptional enzyme molecules akin to “hidden gems” within databases but also enhances the accuracy and precision of the screening process. The emergence of *Kb*PETase provides an improved template for the engineering of high-performance enzymes, expanding the molecular library and offering new tools for the bioconversion and recycling of PET.

## Methods

### Structure search

In the structural search step, we utilized the default setting of FoldSeek server website (https://search.foldseek.com/search). The structure of *Is*PETase (PDB: 5XFY) is selected as the only query protein for FoldSeek search. Target structure database include AlphaFold databases (UniProt50 v4, SwissProt v4, Proteome v4), MGnify-ESM30 (v1), PDB100, and GMGCL 2204.

The core structure is defined by manual structure check of *Is*PETase, ranging from residue 51 to 234, cover the reported catalytic triad and the scaffold of the PET hydrolase. Any searched protein align range not covering this region was excluded from following steps. This structure search step identified 4,235 structural similar protein sequences. Code of this part is available at step 2 in https://github.com/Zuricho/Enzyme-Discovery-Codebase

### Sequence search

In sequence search step, the sequences found in the structure search from different databases were merged and used as queries for the sequence search through the NCBI’s NR database. This step are using MMseqs2 easy-search method with 2 rounds of iterations and set max-seq to 2500, resulting in 33,247,501 sequences. After obtaining this vast repertoire, MMseqs2 easy-cluster method with 50% sequence identity is used to saving the cost for following ProstT5 embeddings computation step. Resulting 436,488 representative proteins derived from 50% sequence identity clusters. Code for this part is available at step 3 and 4 in https://github.com/Zuricho/Enzyme-Discovery-Codebase

### Benchmarking protein language models for protein discovery

We systematically evaluated various computational approaches to assess their effectiveness in identifying and classifying proteins within enzyme commission (EC) groups, in order to find the optimal and most efficient method to discovery novel enzymes. Utilizing a curated dataset of 265,488 protein sequences from the reviewed UniProt database (SwissProt), we focused on proteins up to 1,000 amino acids in length. Our benchmarking compared traditional methods like BLASTp for sequence similarity and FoldSeek with AlphaFold predictions for structural alignment against advanced protein language models such as ESM-2 650M, ESM-1b, and ProseT5. By calculating representation distances and analyzing the area under the ROC curve (AUC) scores for 745 EC groups, we assessed each method’s ability to distinguish between proteins of the same and different EC numbers. This comprehensive evaluation provided insights into the strengths and limitations of each approach, highlighting the potential of protein language models in enhancing protein discovery and classification tasks. Results showed that both the structure-based methods FoldSeek and ProstT5 performed better than BLASTp methods, and ProstT5 is the only unsupervised deep-learning method that performed better than BLASTp (Fig. S6). As a result, we select the structure-aware language model ProstT5 for our further studies. Code for this part is available at https://github.com/Zuricho/protein mining.

### Clustering and representation tree built by structure-aware protein language model

We use protein sequence-to-structure language model ProstT5 to calculate the embeddings of the protein candidates’ sequences. Since ProstT5 is a bi-direction translation model between protein sequence and structure, the “AA2fold” mode of ProstT5 model is used to get a more structure-informed representation based on input sequences. Based on the representation of multiple sequences, we used agglomerative clustering to build a dendrogram for the sequences. The distances between representations are calculated as the Euclidean distance, and the agglomerative clustering method is using ward algorithm from scipy.cluster.hierarchy package. Code of this part is available at step 5 and 6 in https://github.com/Zuricho/Enzyme-Discovery-Codebase.

### Protein expression and solubility prediction

Within the 2 selected cluster in representation tree, there are 6,763 sequences advanced to the next screening step. We used a fine-tuned version of the ESM2^45^ to predict the melting temperature (T_m_) of the candidates using datasets from the literature^46^. Additionally, we employed ProtSolM to predict protein solubility^47^. Only candidates with predicted T_m_ and solubility values exceeding those of the query sequence were retained, narrowing the pool to 223 sequences.

### Protein structure prediction and selection for APET

For further refinement, the structures of the 223 sequences were predicted using AlphaFold2^48^, and any sequence with a pLDDT score below 75 was discarded. Structural alignment was then performed to ensure that the candidate proteins possessed the catalytic active site necessary for PET hydrolase activity. Finally, the candidates were ranked by their predicted T_m_ values, and the top 34 sequences were selected for wet-lab validation.

### Plasmid construction

Genes encoding APETase1-34 were synthesized and optimizated for expression in *Escherichia coli* (Sangon Biotech (Shanghai, China).). The signal peptides of APETase1-34 were predicted with SignalP and removed from the synthetic DNA. The synthesized genes for APETase1-34were cloned into the *Nde*I and *Not*I sites of the pET-28a (+) vector (containing a N-terminal His-tag). A list of nucleotide sequences is provided in Supplementary Table 1.

### Protein Expression and Purification

The expression plasmid was introduced into *Escherichia coli* BL21(DE3) competent cells. A 30 mL seed culture was grown at 37°C in LB medium supplemented with 50 µg/mL kanamycin, and then transferred to a 500 mL shaker flask containing the same antibiotic concentration. The cultures were incubated at 37°C until the OD600 reached 1.0, at which point protein expression was induced by the addition of isopropyl-β-D-thiogalactopyranoside (IPTG) to a final concentration of 0.4 mM. Induction was followed by incubation at 16°C for 16-20 hours. Cells were harvested by centrifugation at 4,000 rpm for 30 minutes, and the resulting pellets were collected for subsequent purification. The cell pellets were resuspended in lysis buffer (25 mM Tris-HCl, 500 mM NaCl, pH 7.4) and disrupted by ultrasonication (Scientz, China). The lysates were then centrifuged at 12,000 rpm for 30 minutes at 4°C, and the supernatants were subjected to Ni-NTA affinity chromatography using an elution buffer (25 mM Tris-HCl, 500 mM NaCl, 250 mM imidazole, pH 7.4). The protein was further desalted with lysis buffer (25 mM Tris-HCl, 500 mM NaCl, pH 7.4) by ultrafiltration. Finally, the fractions containing the purified protein were flash-frozen at -20°C in storage buffer (25 mM Tris-HCl, pH 7.4, 500 mM NaCl, 10% glycerol).

### Differential Scanning Fluorimetry (DSF)

The T_m_ values were determined using the Differential Scanning Fluorimetry (DSF) method with the Protein Thermal Shift Dye Kit (Thermo Fisher, U.S.A). To prepare the reaction mixture, 1.0 μL of SYPRO Orange Dye (SUPELCO, U.S.A) was added to 49 μL of lysis buffer (25 mM Tris-HCl, 500 mM NaCl, pH 7.4). Next, 1 μL of the diluted dye was mixed with 19 μL of protein solution at a concentration of 0.1 mg/mL. DSF experiments were conducted using the LightCycler 480 Instrument II (Roche, U.S.A). The reaction mixture was first equilibrated at 25 °C, then gradually heated to 99 °C at a rate of 0.05 °C/s, where it was held for 2 minutes. Data processing was performed using the Protein Thermal Shift software.

### Nano differential scanning calorimetry (nanoDSC)

nanoDSC measurements were performed by using Nano DSC instruments (TA, U.S.A). The concentration of PETases was 0.5 mg/ml in a buffer containing 25 mM Tris–HCl (7.5) and 500 mM NaCl. All the experiments were carried out at temperatures ranging from 10 to 110°C with a heating rate of 1°C/min and under a pressure of 3 atm. The melting curves of PETases were subtracted from the buffer scans.

### Screening for activity on 4-Nitrophenol butyrate (*p*NPB)

All reactions were conducted in 96-well plates with a total reaction volume of 100 μL. The final concentration of *p*NPB was 0.8 mM, and the enzyme concentration was 100 µg/mL. Each reaction mixture contained 10 μL of *p*NPB solution (dissolved in anhydrous ethanol), 80 μL of 10 mM potassium phosphate buffer (pH 8.0), and 10 μL of enzyme solution, which were incubated at 37°C for 10 minutes. The reaction was terminated by adding 100 μL of anhydrous ethanol to quench the enzymatic activity. Each experiment was performed in triplicate. The concentration of p-nitrophenol was measured at 410 nm using a MD SpectraMax iD5 microplate reader (Molecular Devices, U.S.A.). Enzyme activity was determined by comparing the product concentration to a standard curve of p-nitrophenol (Supplementary Fig. 1).

### PET depolymerization assay using amorphous PET film as substrate

Amorphous PET film (Goodfellow, England) was cut into circular discs with a diameter of 6 mm for each reaction. The discs were incubated in 2900 µL of glycine-NaOH buffer (pH 9.0, 50 mM) containing 100 µL of enzyme solution (stock concentration 0.5 mg/mL) at 30°C, 40°C, 50°C, 55°C, and 60°C for 72 hours. The reaction was terminated by heating the mixture at 90°C for 10 minutes. Each sample was diluted to fall within the linear detection range for terephthalic acid (TPA) and mono(2-hydroxyethyl) terephthalate (MHET). After filtration through a 0.22 µm filter, the assay solution was analyzed by UPLC. All experiments were performed in triplicate.

### UPLC analysis

UPLC analysis was conducted using a Waters ACQUITY Arc system (Waters, U.S.A.) equipped with an autosampler and a UV detector set to 260 nm. The separation was performed on a Kinetex XB-C18 100Å, 5 µm, 50×2.1 mm LC column (Phenomenex, U.S.A.) using a stepped, isocratic solvent gradient. Mobile phase A consisted of water with 0.1% formic acid, and mobile phase B was acetonitrile, with a fixed flow rate of 1.1 mL/min. Samples were injected at either 1 µL or 4 µL. Following injection, the mobile phase was maintained at 13% buffer B for 52 seconds to separate mono(2-hydroxyethyl) terephthalate (MHET) and terephthalic acid (TPA), then ramped up to 95% buffer B for 33 seconds to separate larger reaction products and contaminants. The buffer was then returned to 13% for column re-equilibration, with a total run time of 1.8 minutes. Peaks were identified by comparison to chemical standards prepared from commercial TPA and in-house synthesized MHET, and the peak areas were integrated using software. Under these conditions, TPA eluted around 1.0 minutes, MHET around 2.3 minutes, and small amounts of bis(2-hydroxyethyl) terephthalate (BHET) and longer oligomers eluted between 2.7 and 3.2 minutes. The concentrations of TPA and MHET were determined by constructing standard curves.

### Kinetics analysis

Enzymatic reactions involving 4-nitrophenyl butyrate (*p*NPB) were conducted in 96-well plates with a total reaction volume of 100 μL. The final *p*NPB concentrations ranged from 0.2 to 1.4 mM, while the enzyme concentration was maintained at 0.5 µg/mL. Each reaction mixture contained 10 μL of *p*NPB solution (dissolved in anhydrous ethanol), 80 μL of 10 mM potassium phosphate buffer (pH 8.0), and 10 μL of enzyme preparation, which was incubated at 50°C for 3 minutes. To terminate the reaction, 100 μL of anhydrous ethanol was added. All assays were performed in triplicate. The concentration of p-nitrophenol, the product of the reaction, was measured at 410 nm using a SpectraMax iD5 microplate reader (Molecular Devices, U.S.A.). One unit of enzyme activity was defined as the amount of enzyme required to convert 1 μmol of pNPB per minute. Data analysis was carried out using GraphPad.

### Crystallization and structure determination of *Kb*PETase

Crystals of *Kb*PETase, grown using hanging drop vapor diffusion by mixing equal volumes of protein and a buffer solution containing 1.26 M Sodium phosphate monobasic monohydrate,0.14 M Potassium phosphate dibasic at temperature 291 K. Crystals were rapidly soaked in the reservoir solution supplemented with 20% glycerol as cryo-protectant, mounted on loops, and flash-cooled at 100 K in a nitrogen gas cryo-stream. Crystals Diffraction data was collected from a single crystal at Shanghai Synchrotron Radiation Facility (SSRF) BL18U beamline, China, with a wavelength of 1.75 Å at 100 K. The diffraction data were processed and scaled with HKL-3000 (1). Relevant statistics were summarized in Supplementary Table 3.

The structure was solved by the molecular replacement method with starting model predicted by AlphaFold II. Initial model was build using PHENIX.autobuild (3). Manual adjustment of the model was carried out using the program COOT (4) and the models were refined by PHENIX.refinement (3) and Refmac5 (5). Stereochemical quality of the structures was checked by using PROCHECK (6). All of residues locate in the favored and allowed region and none in the disallowed region. Refinement resulted in a model with excellent refinement statistics and geometry. The structure of *Kb*PETase was deposited in the Protein Data Bank, with PDB code 9IW9.

### Molecular dynamics simulations

The structures of proteins were obtained from PDB database. Protein and a large number of water molecules were filled in a cubic box. Chlorine counter ions were added to keep the system neutral in charge. The CHARMM27 force field was used for the complex and the CHARMM-modified TIP3P model was chosen for water^56-58^. The simulations were carried out at 310 K. After the 4000-step energy-minimization procedure, the systems were heated and equilibrated for 100 ps in the NVT ensemble and 500 ps in the NPT ensemble. The 100 ns production simulations were carried out at 1 atm with the proper periodic boundary condition, and the integration step was set to 2 fs. The covalent bonds with hydrogen atoms were constrained by the LINCS algorithm. Lennard-Jones interactions were truncated at 12 Å with a force-switching function from 10 to 12 Å. The electrostatic interactions were calculated using the particle mesh Ewald method with a cutoff of 12 Å on an approximately 1 Å grid with a fourth-order spline. The temperature and pressure of the system were controlled by the velocity rescaling thermostat and the Parrinello-Rahman algorithm, respectively. All MD simulations were performed using GROMACS 2020.4 software packages.

## Data availability

The refined model of *Kb*PETase generated in this study has been deposited in Protein Data Bank with PDB codes 9IW9. The raw data for figures generated in this study are provided in the Source Data file. The detailed data of all of the PETase discovered can be found in the supplementary data.

## Code availability

The code of the PETase discovery pipeline can be found in https://github.com/Zuricho/Enzyme-Discovery-Codebase.

## Acknowledgements

This work was supported by the grants from the National Science Foundation of China (Grant Number 12104295), the Computational Biology Key Program of Shanghai Science and Technology Commission (23JS1400600), Shanghai Jiao Tong University Scientific and Technological Innovation Funds (21X010200843), and Science and Technology Innovation Key R&D Program of Chongqing (CSTB2022TIAD-STX0017), the Student Innovation Center at Shanghai Jiao Tong University, and Shanghai Artificial Intelligence Laboratory. We would like to thank Dr Jing Wang from Instrumental Analysis Center of Shanghai Jiao Tong University for assistance with PET hydrolysis performance tests via UPLC. Part of the computations in this paper were run on the Siyuan-1 cluster supported by the Center for High Performance Computing at Shanghai Jiao Tong University.

## Author contributions

P.T., L.Z and L.H. conceptualized and supervised this research project. B.Z. and P.T. developed the methodology and designed the pipeline. B.Z., S.J., and M.L. implemented the method and conducted the in-silico screening of PETase. B.W. and R.H. conducted the wet-lab experiments. L.Z. conducted MD simulations. B.W., L.Z., P.T, B.Z., and L.H. wrote the manuscript. All authors reviewed and accepted the manuscript.

